# Across Thermal Scales: Quantized Thermodynamics, Nested Resonant Structures, and the Entropic Stability of Life

**DOI:** 10.1101/2024.02.15.580422

**Authors:** Josh E. Baker

## Abstract

Biological systems are fundamentally containers of thermally fluctuating atoms that through unknown mechanisms are structurally layered across many thermal scales from atoms to amino acids to primary, secondary, and tertiary structures to functional proteins to functional macromolecular assemblies and up. Understanding how chemical thermodynamics scales across these layered structures is central to describing biological structure and function. Muscle – with chemical thermodynamics well-defined on two different thermal scales – provides a clear solution to this problem. In 1938, A.V. Hill made the extraordinary observation that the mechanics and chemistry of muscle contraction is defined independent of the structural components of muscle, implying that the mechanics and chemistry of myosin motor proteins within muscle do not classically scale up to muscle. We have demonstrated experimentally that thermal scaling bridges Hill’s top-down thermodynamics and the bottom-up philosophy of molecular biologists. That is, with thermal scaling *N* individual myosin motor switches physically collapse into an ensemble of *N* myosin motor switches creating a functional entropy within the ensemble well defined by the statistical mechanics of a binary system of switches. This ensemble entropy stabilizes the resonant ensemble structure of muscle and energetically drives the irreversible kinetics and energetics of muscle contraction. Here I develop a general model of thermal scaling and show that it occurs when the ensemble state of a system is defined, at which point the number of ways constituent molecules can account for the ensemble state contributes to the entropy of the ensemble state. At that point, the statistical occupancy of molecular states physically replaces the physical occupancy of molecular states. This is not a classical mechanism and as shown here results in many quantum-like phenomena, which consistent with Hill’s observation means that biological function cannot be described by classical molecular mechanisms.

## I. INTRODUCTION

In 1824, Carnot showed that on account of entropy the mechanical state of a gas cylinder is not defined by the mechanics of individual gas molecules within that cylinder [1]. Similarly, in 1938 A.V. Hill demonstrated experimentally that the mechanical state of muscle is not defined by the mechanics of individual molecules within muscle [2]. A.V. Hill’s results imply that Carnot’s thermodynamics applies to complex biological systems across all thermal scales below that of biological function, which is to say that the function of a biological system is not defined by the function of the constituents (e.g., proteins) of that system. This extraordinary result remains widely ignored because it is unfathomable to most molecular biologists that protein structure-function, which can be precisely measured within individual proteins, does not classically scale up to the structure and function of biological systems defined on larger thermal scales [3–7]. Indeed, one of the most important problems in biology (and physics and chemistry) today remains that of determining how Carnot and Hill’s top-down physics converges with the bottom-up philosophy of molecular biologists [8].

This problem has been solved through experimental studies of muscle contraction [9–12]. Myosin motors in muscle function as force-generating switches induced by actin binding and gated by the release of inorganic phosphate, P_i_ [10,12]. The mechanochemistry of this molecular mechanism is well established through measurements of muscle structure-function, single myosin motor mechanics, and myosin motor structures [10,12–14]. Framed by muscle, the problem then becomes: how does the well-defined mechanochemistry of myosin switches scale up to Hill’s well-defined mechanochemistry of muscle? Rephrased, what is the physical difference between *N* individual myosin switches (e.g., observed in *N* different single molecule experiments) and an ensemble of *N* myosin switches (e.g., observed in muscle)? The simple answer is that the latter is a binary system of switches with an ensemble entropy defined by statistical mechanics [15] and the former is not. In other words, thermal scaling creates entropy contained within thermal fluctuations delocalized among motor switches in the form of heat.

The effect of this entropy on muscle mechanochemistry is described by the Gibbs reaction free energy equation (the change in system free energy with the extent of a reaction). Specifically, the actin-myosin binding reaction that induces the switch occurs with a change in state of the binary system, which means that irreversible actin-myosin binding and muscle’s irreversible power stroke are energetically driven by an increase in the ensemble entropy of the binary system of switches [15]. This physical, functional ensemble entropy created by thermal scaling has been directly observed in muscle [9] and the entropically-driven irreversible energetics and kinetics of an ensemble of force-generating myosin switches accurately describe most key aspects of muscle contraction [15–17]. In a companion article, I develop the implications of this irreversible entropically driven chemical reaction for our understanding of both chemical kinetics and the physics of the arrow of time.

Here, I assume that the same entropically-driven thermal scaling occurs across all thermal scales in biological systems from atoms to amino acids to peptide chains to secondary, tertiary and quaternary structures to protein ensembles to large filamentous assemblies to cells to tissues and organs and organisms [14,18–20]. In general, *N* degrees of freedom of an ensemble of *N* structural elements physically collapse to create *N* discrete states of a new resonant structure on a new thermal scale. This collapse occurs when the state of an ensemble system is defined at which point the number of ways molecules can occupy molecular states to account for the ensemble system state contributes energetically to the state of the system as entropy. At this point, the physical occupancy of molecular states (chemical activity) is physically replaced by the statistical occupancy of molecular states (entropy). Ensembles of ensemble structures collapse to create new structures on even larger thermal scales, and this process repeats across all thermal scales, creating resonant structures and useful entropy nested from amino acids to functional biological systems.

Thermal scaling is directly observed in muscle [9], provides the physical basis for Hill’s thermodynamic muscle model [11], and accounts for most key aspects of muscle contraction [8,16,17]. The physics of thermal scaling predicts an observational uncertainty when measuring the states of individual molecules (defined on thermal scale i) within an ensemble (defined on thermal scale j), implying that thermal scaling is quantum-like. Thus, thermal scaling disproves classical mechanisms of biological function [3–7] and demonstrates how life is formed and functions not in local opposition to the second law of thermodynamics but because of it.

## II. THE COLLAPSE OF TWO CHEMICAL STATES INTO ONE THE k_B_T·ln2 PARADOX

Figure 1A illustrates a simple reaction where a structural element equilibrates between one state A and two energetically equivalent states B_1_ and B_2_ with an equilibrium constant *K* for each reversible transition. If differences between A and B but not between B_1_ and B_2_ are measurable, the observed equilibrium constant between A and B (B_1_ or B_2_) is 2*K*. This sets up a Gibbs-like paradox. That is, as B_1_ and B_2_ become increasingly similar, the equilibrium constant remains 2*K* up until B_1_ and B_2_ become identical at which point the equilibrium constant becomes *K*. The paradox is that a relatively large energy change

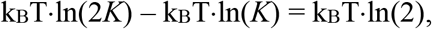

is created by an infinitesimally small change in the physical difference between two states. The solution to this paradox depends on the nature of the difference between B_1_ and B_2_.

**Figure 1.**
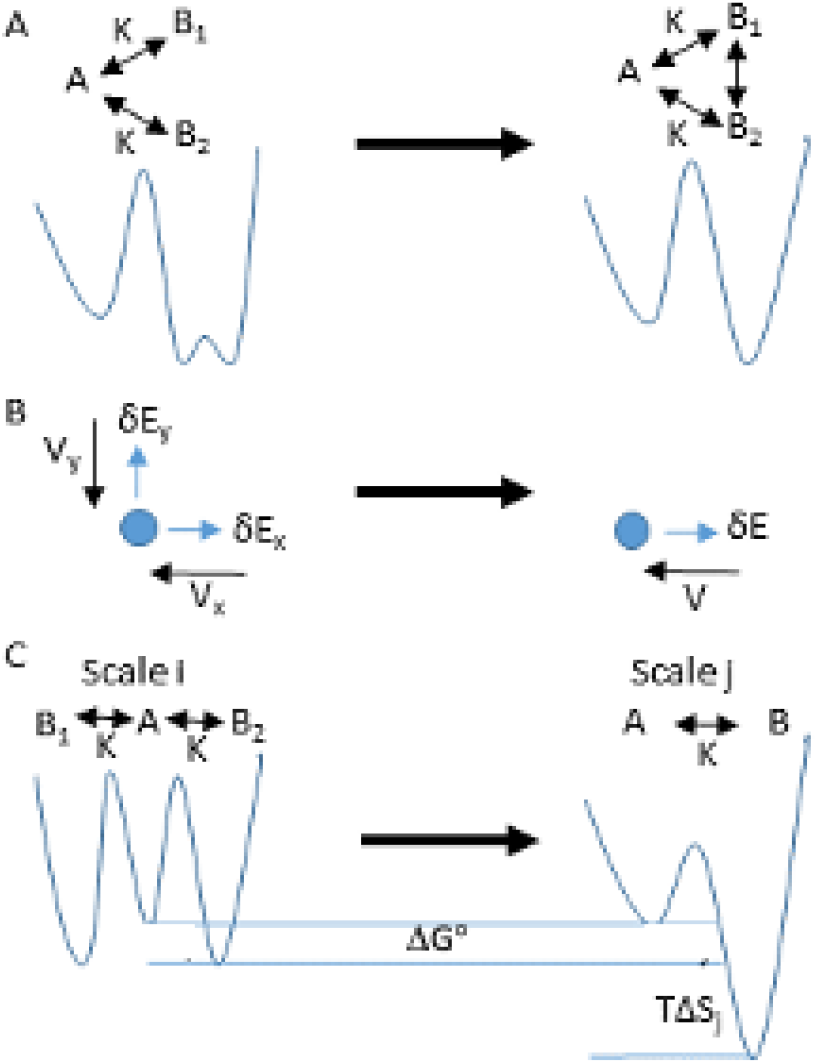
The collapse of two degrees of freedom into one. (A) A three-state kinetic scheme (top) and corresponding energy landscape (bottom) for an equilibrium reaction between states A, B_1_, and B_2_, where B_1_ and B_2_ are energetically equivalent states. If B_1_ and B_2_ are defined within a single degree of freedom they can be coarse-grained together (left to right). (B) A particle thermally fluctuating in two dimensions, x and y, (two degrees of freedom) against a potential *V*_*x*_ and *V*_*y*_ has energy states *dE*_*x*_ and *dE*_*y*_. When these two dimensions collapse into one (left to right) the particle moves in one dimension against a potential, *V*, with an energy *dE*. (C) The kinetic scheme (top) and corresponding energy landscape (bottom) in (A) where B_1_ and B_2_ are now defined as separate degrees of freedom on scale i (left) that collapse into a single degree of freedom, B, on scale j (right).

If B_1_ and B_2_ are differences within a single degree of freedom, B, they can be treated as substates (part of a rugged landscape) of B (Fig. 1A). These substates can be coarse grained (smoothed, Fig. 1A left to right) to describe an equilibrium between A and a coarse-grained B. In this case, as B_1_ and B_2_ become increasingly similar the shape of the coarse-grained energy well, B, changes smoothly with no discrete energy change when B_1_ and B_2_ become equivalent.

However, if B_1_ and B_2_ describe two different degrees of freedom, the paradoxical k_B_T·ln(2) remains problematic because degrees of freedom cannot be smoothly coarse-grained away (e.g., the average of two and one degrees of freedom, 1.5, is not defined).

Figure 1B illustrates a particle moving in one of two dimensions (two degrees of freedom) within two potentials, V_x_(x) and V_y_(y). Starting in a reference state, an infinitesimally small change in energy, δE, can occur with a displacement in either direction, δE_x_ or δE_y_.

Because δE_x_ and δE_y_ are defined by two discretely different degrees of freedom, even infinitesimally small energy changes cannot be smoothly averaged (coarse-grained) together. Still, muscle tells us that multiple degrees of freedom do physically collapse into one (Fig, 1B, left to right). While the two states, δE_x_ and δE_y_, are each occupied in one way (Ω_x_ = Ω_y_ = 1) the collapsed state, δE, can be occupied in two ways (δE_x_ or δE_y_, Ω = 2). Thus, according to Boltzmann’s definition of entropy, S = k_B_·lnΩ, when two degrees of freedom collapse into one, entropy increases by

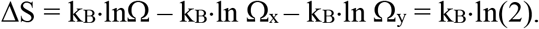

It follows from the second law of thermodynamics (dQ = TdS) that the paradoxical k_B_T·ln(2) is quantized heat, dQ, generated when two degrees of freedom collapse into one. If the change in energy, δE_x_ or δE_y_, defined on the thermal scale of x and y approaches zero and only thermal energy, ½k_B_T, exists for each degree of freedom, the minimum (quantized) δE that can be defined on the collapsed thermal scale (Fig. 1B, right) is the heat, k_B_T·ln(2), delocalized between x and y.

In general, when *N* degrees of freedom collapse into one, thermal energy, (*N*/2)k_B_T, is lost and entropy, k_B_Tln(*N*), and thermal energy, ½ k_B_T, are gained on a new thermal scale. When ½(*N* – 1) > ln(*N*), excess heat is lost with this collapse, which means it is spontaneous and irreversible. This occurs when more than three degrees of freedom collapse into one. We refer to this as thermal scaling. With thermal scaling, the loss of the physical occupancy of states with a gain in statistical occupancy of states results in observational uncertainty, or the inability to define the states of the constituents of a system (their degrees of freedom are no longer defined) on the thermal scale of the system (A.V. Hill’s observation). For example, on the thermal scale of the collapsed state in Fig. 1B, δE_x_ and δE_y_ both physically exist with equal probability at a given instant of time within a resonant structure. To determine δE_x_ and δE_y_ energy is required to reverse the collapsed degrees of freedom.

## III. BOLTZMANN AND GIBBS BOTH ACCOUNT FOR ΔS = k_B_T·ln(2)

The increase in entropy, k_B_T·ln(2), with a collapse of two degrees of freedom is also described by the Gibbs free energy equation. Figure 1C revisits the reaction in Fig. 1A only here B_1_ and B_2_ represent two different degrees of freedom. Assume on the thermal scale of B_1_ and B_2_ (scale i), an equilibrium constant between state A and B_1_ or B_2_ of 2*K*, which is *K* for each degree of freedom. The free energy difference between states A and B_1_ or B_2_ is then ΔG_i_ = –k_B_T·ln(2*K*). On the thermal scale of B (scale j), the equilibrium constant is *K* because only one degree of freedom and one state, B, exist. The free energy difference between states A and B must be the same on thermal scales i and j, and so ΔG_j_ = –k_B_T·ln(2*K*). The entropic contribution to the free energy is ΔS_j_ = k_B_·ln(2), and the enthalpic contribution is the measurable component of the free energy on scale j, which is ΔH_j_ = k_B_T·ln(*K*).

Thus, on thermal scale i

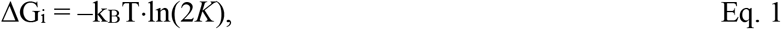

and on thermal scale j,

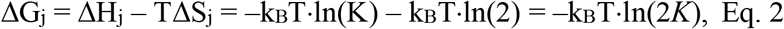

where the observable free energy change for a single degree of freedom, ΔGº = ΔH_j_, defined and measurable on both scales, is the standard reaction free energy.

While mathematically equivalent, these equations describe two physically different systems (i and j). On thermal scale i, k_B_T·ln(2) describes the physical ways that B_1_ and B_2_ can be occupied (either one or the other state is physically occupied). On thermal scale j, k_B_T·ln(2) describes the a priori ways that B can be occupied (it is two, independent of the physical occupancy of B_1_ and B_2_). The latter physically tilts the energy landscape (Fig. 1C, right) to create, on a new thermal scale, a structure B that physically replaces structures B_1_ and B_2_. In a companion article, a similar mathematical equivalence is shown between chemical kinetics defined by chemical activities on scale i (a physical occupancy of states that pushes a reaction forward) and chemical kinetics defined entropically on scale j (a statistical occupancy of states that pulls a reaction forward).

## IV. THE COLLAPSE OF *N* INDIVIDUAL ELEMENTS INTO AN ENSEMBLE OF *N* ELEMENTS

Consider an ensemble of *N*_*i*_ structural elements (for simplicity I assume one degree of freedom per structural element) each with a quantized energy change δ*E*_*i*_. The number of ways these *N*_*i*_ elements can have a total energy *x*_*i*_·δ*E*_*i*_ is Ω_i_ = *N*_*i*_!/[*x*_*i*_!(*N*_*i*_ – *x*_*i*_)!], where *x*_*i*_ is the state of the ensemble. The total number of states, *x*_*i*_, is *N*_*i*_, and the entropy, S_i_(*x*_*i*_) = k_B_·lnΩ_i_, of the ensemble in state *x*_*i*_ is

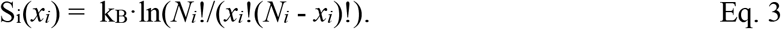

The heat contained by the ensemble in state *x*_*i*_ is then Q_i_(*x*_*i*_) = TS_i_(*x*_*i*_), which is plotted in Fig. 2 for values of *N*_*i*_ of 3, 5, and 20, showing that an ensemble of *N*_*i*_ structural elements collapses into an entropic well that defines the energy landscape of an ensemble structure (the change in energy with a change in state of the ensemble structure). Clearly, when the state of the ensemble system is defined, the statistical occupancy of individual structural elements replaces the physical occupancy of individual structural elements.

**Figure 2.**
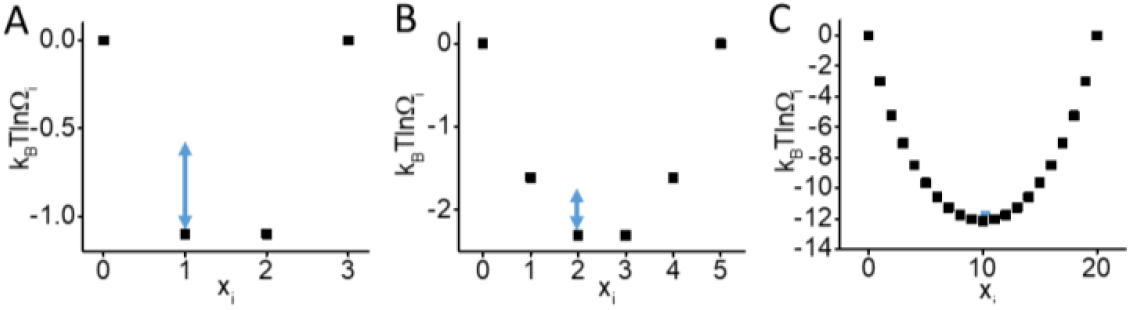
Entropic Wells. (A – C) Equation 3 (k_B_T·lnΩ_i_) is plotted versus *x*_*i*_ for *N*_*i*_ = 3, 5, and 20. The blue double arrow is ½k_B_T.

On this new thermal scale, *N*_*i*_ individual structural elements physically create one new ensemble structure having *N*_*i*_ structural states where the energetics of changes with changes in ensemble chemical state are defined by an entropic reaction energy landscape. Thermal fluctuations within this ensemble landscape (blue double arrow in Fig. 2 indicates ½k_B_T thermal energy) are driven by less thermal energy than that in its *N*_*i*_ structural components because according to equipartition the physical collapse of *N*_*i*_ degrees of freedom into one degree of freedom decreases thermal energy. In an accompanying article, I describe the difference between chemical kinetics described by diffusion within the energy landscapes of *N*_*i*_ individual structural elements and chemical kinetics described by diffusion within the reaction energy landscape of an ensemble of *N*_*i*_ structural elements.

Because it involves the creation of entropy, thermal scaling is physical. Energy is conserved because the entropy gained with thermal scaling is the thermal energy of the structural components transformed into heat (entropy) delocalized among the structural components of the ensemble structure. If the net gain in heat exceeds that which can be delocalized within the ensemble structure, thermal scaling is irreversible, energetically stabilizing the ensemble structure. We have experimentally demonstrated that thermal scaling occurs in muscle when *N* individual molecular switches physically collapse into an ensemble of *N* molecular switches to create muscle’s ensemble (macromolecular) structure with an entropic well defined as in Fig. 2 by the statistical mechanics of a binary system of switches [9]. We have shown that irreversible power output by muscle occurs with an entropically-driven chemical reaction down this well [11,15].

The entropic wells in Fig. 2 resemble entropic funnels in models of protein folding [20]; only here the implication is that protein folding occurs not through a rugged landscape but through nested funnels defined on different thermal scales. For example, secondary structures fold within entropic wells defined by the degrees of freedom of primary structure; tertiary structures fold within entropic wells defined by the degrees of freedom of secondary structures; etc. The formation of each new entropic well at each new thermal scale is irreversible and stabilized if the heat created through thermal scaling exceeds that which can be contained within the ensemble structure (Fig. 1). The energy levels of the resulting entropic well are quantized by the discrete number of degrees of freedom that collapse to create it. Here, I refer to the difference between the first energy level and the energy minimum as the ground state energy spacing for this structure. In muscle, we have shown that resonant structural elements of contractile proteins thermally scale all the way up to muscle’s one degree of freedom [15], which is to say that muscle is energetically stabilized not like a house of cards against the second law of thermodynamics but because of it.

Thermal scaling requires that all *N*_*i*_ structural elements have similar changes, δ*E*_*i*_, in their ground state energy spacing, which is to say all *Ni* structural elements must have similar numbers of structural states (Fig. 3A). This is analogous to resonant frequencies in a normal model analysis. While the entropic wells in Fig. 3A are created from the number of ways an ensemble of structural elements changes by δ*E*_*i*_, excited states can be created from the number of ways an ensemble of structural elements changes by 2δ*E*_*i*_, 3δ*E*_*i*_, etc., (blue lines in Fig. 3A). This is analogous to harmonics in a normal model analysis. For entropic wells created by large *N*_*i*_, energy levels become less quantized and thermal fluctuations are minimized (Fig. 2) at which point thermal scaling reaches a classical limit.

**Figure 3.**
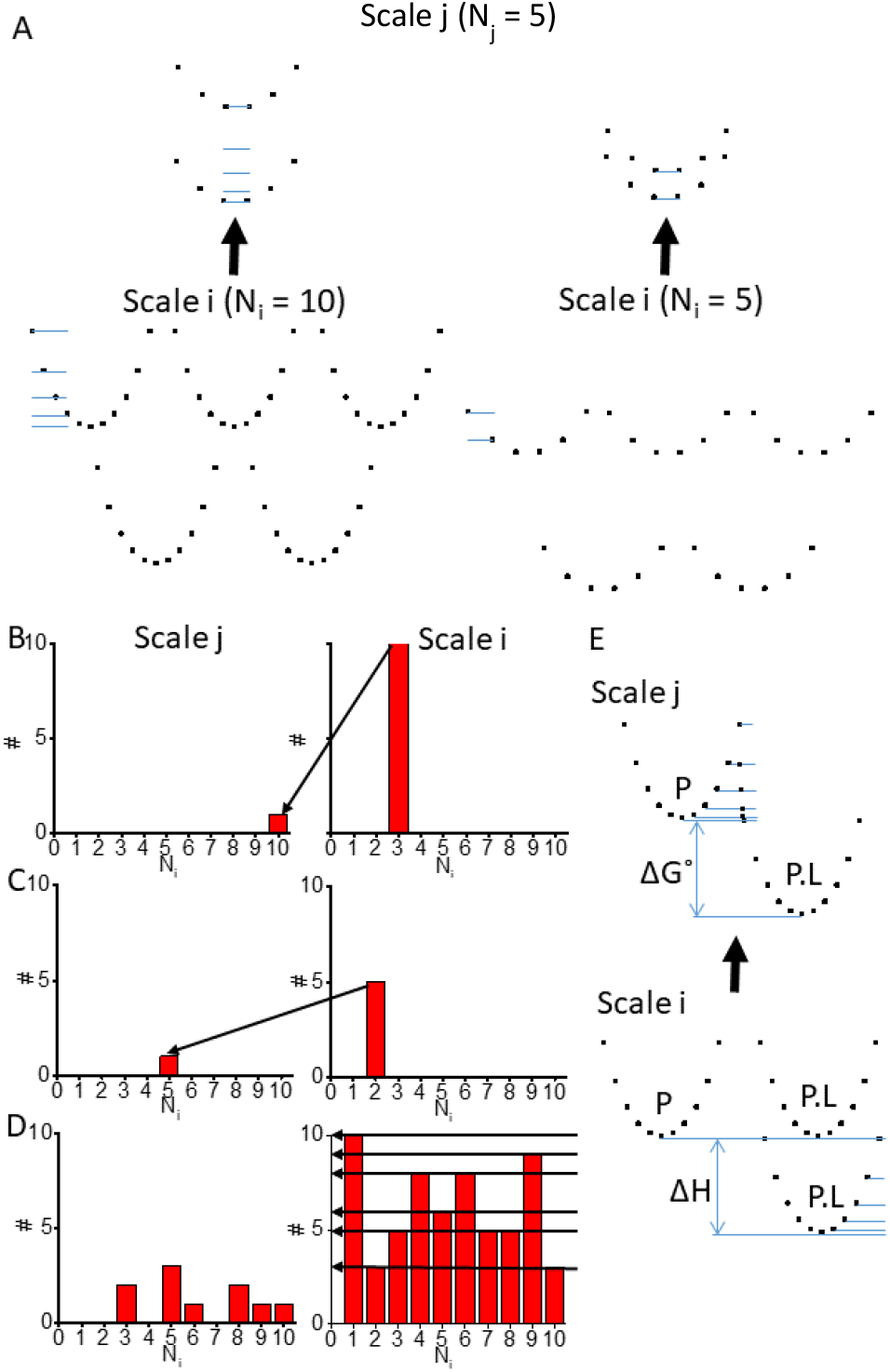
Nested Resonant Structures. (A) Entropic wells (Eq. 3) corresponding to ten ensemble structures on thermal scale i are plotted, five composed of *N*_*i*_ = 10 elements and five composed of *N*_*i*_ = 5 elements. The two sets of five ensemble structures collapse into two new ensemble structure on a higher thermal scale, j (bottom to top) both composed of *N*_*j*_ = 5 elements. The blue lines indicate the excited energy states of wells on scale j created from higher level energy spacings (e.g., 2*dE*, 3*dE*, etc.) on scale i. (B – D) Histograms of hypothetical populations of ensembles of *N*_*i*_ structural elements (x axis) on scale i that collapse (right to left) into ensembles of *N*_*j*_ structural elements (x axis) on scale j. (E) Two sets of ensemble structures (*N*_*i*_ = 10) on scale i, one with ligand bound and the other with no ligand bound are separated energetically by the ligand binding enthalpy, ΔH. An excited state of the ligand bound structures collapse with apo structures to create a resonant tunnel (activation energy barrier) between the ligand bound and apo structures (chemical states).

Histograms of populations of structural elements on scale i with different numbers of states, *N*_*i*_, are illustrated in Figs. 3B – 3D showing how these populations collapse into new structures on scale j. At one extreme, Fig 3B describes a monatomic gas of *N* molecules with three degrees of freedom. Here, the total, 3^N^, degrees of freedom collapse into one degree of freedom, generating *N*·k_B_T·ln(3) of quantized heat. This heat is not contained within entropic structural wells because there are no bonds between atoms to contain it. Instead, it is contained by the walls of a cylinder and thus is balanced against the pressure, P, and volume, V, of the cylinder as PV = N·k_B_T·ln(3), resembling the ideal gas law [ln(3) ≅ 1.1].

As another example, *N* two-state switches connected in series through flexible linkages (Fig. 3C) collapse into a single entropic well (a polymer) with *N* states. This energy well describes the potential energy landscape (Fig. 2) of an entropic spring [15]. There are many intermediate resonant structures that must be created between the thermal scales of atoms and muscle, and Fig. 3D illustrates how populations of different structural elements on thermal scale i collapse to form a different set of structures on thermal scale j. It is clear from Fig. 3D that large structures can contain quantized heat if structural elements are incrementally layered across many thermal scales, which is the structural layering that has evolved in biological systems.

## V. LIGAND BINDING: THE COLLAPSE OF TWO STRUCTURES WITH AND WITHOUT BOUND LIGAND

Figure 3E illustrates two different structural states (i.e., chemical states) on scale i corresponding to a structure with and without a bound ligand. The two structural elements differ in energy by the molecular binding free energy (Fig. 3B), and this difference propagates across all scales (Eqs. 1 and 2). If excited states of the ligand-bound structure (Fig. 3A) overcome the free energy difference between chemical states, structural elements from both chemical states collapse together (Fig. 3E, bottom to top) to create an entangled structure on scale j. The resulting activation energy barrier between the two states is a resonant structure defined by thermal energy delocalized among structural elements both with and without bound ligand. Thus, state transitions occur through quantum-like tunnels between states with structural details entangled across thermal scales (Fig. 3E). In other words, state transitions are described by quantum-like entanglement not the classical mechanics of Eyring and Kramers [4,21].

## VI. APPLICATION TO MUSCLE

As illustrated in Fig. 4, in muscle *N* individual force-generating myosin motor switches (Fig. 4A) thermally scale up to an ensemble of *N* myosin switches with *N* chemical states (Fig. 4B). Individual myosin switches are induced by actin binding, which is shown in Fig. 4A (top) as a chemical transition from state A to B. For an individual myosin switch, the energy landscape for an A to B transition is illustrated in Fig. 4A (bottom), where here I arbitrarily assume that each myosin motor is an ensemble of 11 structural components. The free energy difference between states A and B is ΔGº. When 20 individual myosin motor switches thermally scale up to an ensemble of 20 myosin motor switches, the ensemble energy landscape of muscle is created on a higher thermal scale (Fig. 4B, black squares) with 20 states.

**Figure 4.**
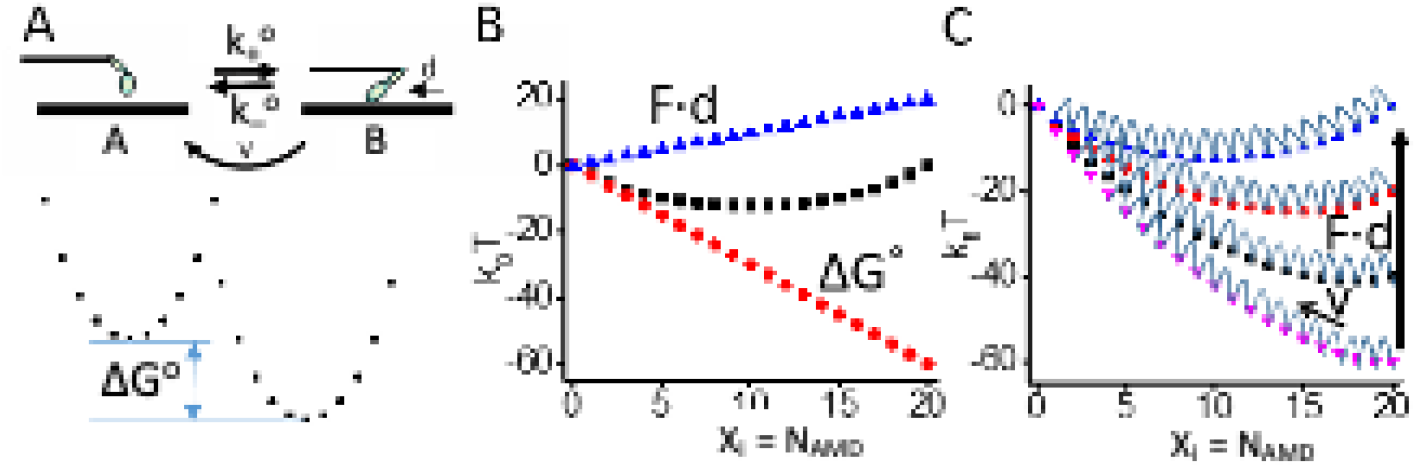
Thermal scaling of muscle mechanochemistry.(A) A myosin motor (M, ovals) is a protein switch induced by actin (A, thick black line) binding and gated by phosphate (P_i_) release with a throw of *d*. The switch is a reversible intermediate step in the actin-myosin catalyzed ATP hydrolysis reaction, *v*, with forward and reverse rates, *f*_*+*_ and *f*_*–*._ The entropic wells for the protein switch (energy landscape of chemical reaction) illustrate the enthalpic binding energy, ΔH (bottom) (B) As described in [15], the entropic well (Eq. 3) created when *N*_*j*_ = 20 myosin motors collapse into a macromolecular actin-myosin assembly is plotted (black symbols) as a function of the extent of the reaction, *x*_*i*_, or the number, *N*_*AMD*_, of myosin motors in state AMD. The free energy for the macromolecular complex decreases by ΔGº with each chemical step (red symbols). The internal work performed on the system by a chemical step increases linearly with the extent of the reaction with a slope proportional to muscle force, *F* (blue symbols). (C) The sum of the energetic contributions to the macromolecular landscape are plotted for different values of F illustrating the activation energy barrier for each step.

In addition to entropy, both enthalpy and non-PV work contribute to muscle’s ensemble free energy landscape. On the thermal scale of muscle, the free energy of the ensemble system increases by ΔGº with each chemical step (Fig. 4B, blue triangles). As shown by A.V. Hill [2] and our research group [15], non-PV work varies linearly with muscle force, F, and adds to the free energy of the ensemble system with each chemical step (Fig. 4B, red circles). The offsetting effects of ΔGº and *F* in Fig. 4B have been observed in muscle [9]. The free energy landscape of the ensemble muscle system is the sum of these energetic components (Fig. 4C) and is the landscape within which muscle contracts. The catalyzed ATPase reaction, *v*, does not affect the landscape, rather it transfers myosin motors from B to A within the landscape (arrow, *v*). If muscle is allowed to shorten, the chemical relaxation back toward an energy minimum performs external work, which diminishes the contribution of ΔGº to the landscape. If muscle is held at a fixed length, the chemical relaxation generates muscle force, *F*, which changes the landscape as illustrated in Fig. 4C. Perturbations to small scale structural elements (mutations, ligand binding, post-translational modifications, etc.) affect muscle mechanics through these thermodynamic parameters not through molecular mechanisms. This simple model accurately describes most key aspects of muscle contraction [8,15–17].

## VII. DISCUSSION

Muscle evolved, not by following the rules of science as we understand them today but by following universal physical laws that have existed for billions of years. It is therefore no surprise that muscle has taught us new perspectives on these physical laws. A.V. Hill showed that muscle force is not determined by the mechanical states of individual myosin motors contained within it [1], yet we observe that individual myosin motors within muscle generate force [10,13,22]. Clearly, the bottom-up philosophy of molecular biologists does not classically scale up to the top-down physics of Hill’s thermodynamics. From the penultimate scale to the thermal scale of muscle contraction we have observed that *N* individual force-generating myosin motor switches on thermal scale i physically collapse into an ensemble of *N* switches on thermal scale j creating entropy within the ensemble (the entropy of a binary system of switches) [9,15]. The state of this binary system changes with actin-myosin binding (a flip of a switch), which means that actin-myosin binding and muscle’s power stroke are irreversibly driven by a change in the entropy of binary system of switches defined on scale j. Here, we extend this model to describe thermal scaling of *N* molecules on scale i to an ensemble of *N* molecules on scale j that I propose occurs across every thermal scale in biology from amino acids to primary, secondary, tertiary, and quaternary protein structure to protein assemblies on up. In a companion article, I describe the difference between chemical kinetics defined on scales i and j.

The creation of entropy upon thermal scaling occurs when the state of the ensemble is defined, at which point the number of ways the constituents of the ensemble can account for the ensemble state contributes to the energy of the ensemble state as entropy. With this physical transition, the physical occupancy of molecular states is replaced by the statistical occupancy of molecular states, leading to quantum-like phenomena. As shown here, thermal scaling creates resonant structures, predicts observational uncertainty when measuring the state of individual molecules in state i from an ensemble state j, and implies that chemical state transitions occur with tunneling through component structures entangled by thermal scaling. Thermal scaling explains Schoedinger’s cat (across scales of light) and Maxwell’s demon (across scales of thermal energy); it provides a solution to the Gibbs paradox [23]; it suggests an entropic interpretation of the ideal gas law; and it implies that life on earth is energetically stabilized not against the second law of thermodynamics but because of it.

Two different entropically driven processes are important in this analysis. Here, I describe an irreversible creation of ensemble entropy with increasing thermal scale that energetically stabilizes resonant ensemble structures. In a companion article, I describe how within a thermal scale a change in state of the ensemble structure occurs with an entropically driven irreversible chemical reaction (the arrow of time). We have shown that this thermodynamic formulation of chemical kinetics accurately describes steady state and transient muscle mechanics, kinetics, and energetics [11,15].

Thermal scaling resembles a normal mode analysis [24] of protein dynamics where different vibrational modes with different thermal distances occur on different thermal scales. However, here the vibrations are thermal fluctuations not harmonic oscillations, and the “modes” are quantized energy spacings not frequencies. Moreover, the transition between scales is an irreversible, physical transformation that creates ensemble entropy that is physically stored within ensemble structures; it is not an increase in entropy with coarse graining described as a loss of information associated with a mental state of not knowing.

On the thermal scale of atoms, the largest ground state energy spacing is k_B_T·ln(2), setting an upper limit for the energy that can be absorbed by a resonant structure created when two degrees of freedom collapse into one. Thermal energy is quantized by this spacing; however, quantized light, hv, can also be absorbed by this spacing at a frequency defined by k_B_T·ln(2) = h*v*, or *v* = 200 cm^-1^ at room temperature. This is the highest energy of quantized light that can be absorbed by a structural element as quantized heat. It is commonly assumed that structural elements absorb infrared light because its frequency, *v*, matches the frequency of harmonic oscillations of structural elements. Here, structural elements absorb infrared light because its energy, hv, matches the quantum heat spacing of resonant structural elements. This is the bridge from quantized light to quantized heat, which then spans across many thermal scales all the way up to the thermodynamics of muscle contraction.

Muscle shines a light on how the world works across all thermal scales within which it evolved and now functions. The collapse of degrees of freedom creates entropic wells within which proteins fold; large aggregate structures are formed; and muscle contracts. Extended to larger scales, it is a reasonable hypothesis that a collapse of degrees of freedom creates entropic wells (e.g., gravitational potentials) that stabilize the aggregation of masses [25]. In general, a change in our understanding of scaling (over energy scales of light, heat, or larger scale fluctuations) from that of a smooth, reversible, mental exercise to that of a discrete physical transformation provides new perspectives on phenomena ranging from the atomic to cosmological.

